# Skeletal muscle-derived extracellular vesicles are altered with chronic contractile activity: a dynamic light scattering analysis

**DOI:** 10.1101/2022.02.25.481852

**Authors:** Patience O. Obi, Emily Turner-Brannen, Adrian R. West, Hagar I. Labouta, Joseph W. Gordon, Ayesha Saleem

**Author notes:** To whom correspondence should be addressed: **Ayesha Saleem, PhD**, Associate Professor, Faculty of Kinesiology and Recreation Management University of Manitoba, Winnipeg, MB, R3T 2N2, 120 Frank Kennedy Centre | |, Research Scientist, Children’s Hospital Research Institute of Manitoba (CHRIM) John Buhler Research Centre (JBRC), Winnipeg, MB R3E 3P4, 600A - 715 McDermot Avenue | |.

## Abstract

Extracellular vesicles (EVs) are small lipid bilayer-delimited particles that are secreted by all cells, playing a central role in cellular communication. EVs are released from skeletal muscle during exercise, but the effects of contractile activity on skeletal muscle-derived EVs (Skm-EVs) are poorly understood due to the challenges in distinguishing Skm-EVs derived from exercising muscle in vivo. Using tunable resistive pulse sensing (TRPS), we previously demonstrated that chronic contractile activity (CCA) increased the secretion of Skm-EVs from C2C12 myotubes, while their size and zeta potential remained unchanged. Here, we aimed to determine whether similar results could be obtained using an alternative method of EV characterization, dynamic light scattering (DLS). C2C12 myoblasts were differentiated into myotubes, and electrically paced (3h/day x 4days @14V, C-PACE EM, IonOptix) to mimic chronic exercise in vitro. EVs were isolated from conditioned media of control and stimulated myotubes using differential ultracentrifugation, and characterized based on size and zeta potential. The mean size of EVs from chronically stimulated myotubes (CCA-EVs, 132 nm) was 26% smaller than control (CON-EVs, 178 nm). Size distribution analysis revealed that CCA-EVs were enriched in small EVs (100-150 nm), while CON-EVs were largely composed of 200-250 nm sized vesicles. Additionally, zeta potential was 27% lower in CCA-EVs compared to CON-EVs. Our data indicate that the effect of CCA on facilitating the release of smaller, more stable EVs, is a robust finding, reproducible by multiple methods of EV characterization. Future studies investigating the mechanisms by which CCA influences Skm-EV biogenesis and secretion are warranted.

## Introduction

Extracellular vesicles (EVs) are small, lipid bilayer-enclosed particles that facilitate intercellular communication by transferring bioactive molecules such as proteins, lipids, and nucleic acids [1]. Traditionally, EVs have been categorized into three subtypes based on their origin, size, and biological characteristics. These include exosomes (30–150 nm), which originate from the inward budding of endosomal membranes, forming intraluminal vesicles within multivesicular bodies (MVBs) that are later released via exocytosis; microvesicles (100–1000 nm), which arise from the direct outward budding of the plasma membrane; and apoptotic bodies (500–5000 nm), which are shed from the membrane of apoptotic cells [2]. According to the Minimal Information for Studies of EVs (MISEV) guidelines, EVs can also be classified based on size—where small EVs (sEVs) are <200 nm and large EVs (lEVs) exceed 200 nm— biochemical composition, and cell of origin [3].

Skeletal muscle is a secretory organ, releasing EVs that may influence both local and systemic adaptations to exercise [4], [5]. We have previously demonstrated that skeletal muscle-derived EVs (Skm-EVs) released following chronic contractile activity (CCA) promote mitochondrial biogenesis in recipient C2C12 myoblasts [6]. Additionally, we demonstrated that these CCA-EVs reduced cell growth and increased cell death in lung cancer cells [7], suggesting that muscle-derived EVs may have important implications beyond muscle adaptation, including potential roles in cancer biology. Understanding how contractile activity alters Skm-EV biogenesis and characteristics is therefore essential to elucidating their functional impact.

Previously, using an orthogonal, single-particle tunable resistive pulse sensing (TRPS) analysis of Skm-EVs, we observed an increase in EV secretion following CCA, while their size and zeta potential remained unchanged [6]. However, variations in EV characterization methods can yield different outcomes [8], [9]. Thus, using complimentary EV characterization methodologies is recommended to bolster findings. Dynamic light scattering (DLS) is an alternative technique that provides insights into the size distribution and surface charge of EVs, offering a complementary approach to TRPS [10], and has been used previously by our group [11], [12]. The extent to which DLS produces findings consistent with TRPS in the context of Skm-EVs remains unclear. In this study, we aimed to characterize Skm-EVs isolated from electrically stimulated C2C12 myotubes using DLS and compare our findings to previous TRPS-based analyses.

## Methods

### Cell culture

Cell culture was perfomed as previously described [6]. Murine C2C12 myoblasts were seeded in a six well plate pre-coated with 0.2% gelatin, and grown in fresh Dulbecco’s Modification of Eagle’s Medium (DMEM; Sigma-Aldrich) supplemented with 10% fetal bovine serum (FBS; Gibco/ThermoFisher Scientific) and 1% penicillin/streptomycin (P/S) (growth media). Cells were grown at 37 °C in 5% CO2 incubator for 24h. When myoblasts reached approximately 90-95% confluency, the growth media was switched to differentiation media (DMEM supplemented with 5% heat-inactivated horse serum (HI-HS; Gibco/ThermoFisher Scientific) and 1% P/S) for 5 days to differentiate myoblasts into immature muscle fibers, or myotubes.

### Chronic contractile activity (CCA) and and sample collection for EV isolation

CCA was performed as previously described by our group [6]. After 5 days of differentiation, myotubes were divided into control (CON) and CCA groups. The CCA plates were stimulated using the C-Pace EM cell culture stimulator, with C-dish and carbon electrodes (IonOptix, Milton, MA, United States) while the CON plates had the C-dish placed on top but without the carbon electrodes. Myotubes were subjected to CCA at 1 Hz (2 ms pulses), 14 V for 3 h/day for 4 consecutive days. After each bout of contractile activity, spent media was discarded and 2 mL fresh differentiation media added to each well. This is important as previous observations by Saleem and Hood (unpublished, PhD dissertation) indicate that changing spent media post-contractile activity improves protein and RNA yield from CCA myotubes. Myotubes were allowed to recover for 21 h before the next bout of contractile activity. On day 4, immediately after the last bout of contractile activity, media was changed to sEV-depleted differentiation media (DMEM supplemented with 5% sEV-depleted HI-HS and 1% P/S) for both groups, with an additional 21 h recovery period. sEV-depleted differentiation media was used to remove any confounding effects of EVs found naturally in horse serum. After 4 days of CCA, post-21 h recovery, 12 mL conditioned media from CON and CCA myotubes was collected and used for EV isolation and characterization as described below.

### Isolation of EVs by differential centrifugation

EVs were isolated using dUC as previously described [6]. Briefly, conditioned media (12 mL per six-well plate) was collected and sequentially centrifuged at 300 × g for 10 min at 4 °C (Sorvall™ RC 6 Plus Centrifuge, F13-14 fixed-angle rotor) to remove dead cells, followed by 2000 × g for 10 min at 4 °C to eliminate cell debris. The supernatant was then centrifuged at 10,000 × g for 30 min at 4 °C to remove large vesicles. To obtain the sEV-enriched fraction, the remaining supernatant was ultracentrifuged at 100,000 × g for 70 min at 4 °C using a Sorvall™ MTX 150 Micro-Ultracentrifuge (S58-A fixed-angle rotor). The resulting pellet was resuspended in 1 mL PBS and subjected to a second ultracentrifugation at 100,000 × g for 70 min at 4 °C. The final sEV pellet was resuspended in 50 µL PBS and used for subsequent analysis.

### Dynamic light scattering (DLS)

DLS analysis of EVs was performed as previously described [11]. Measurements of EV size and zeta potential were performed by phase analysis DLS using a NanoBrook ZetaPALS (Brookhaven Instruments, Holtsville, NY, USA) in collaboration with Dr. Hagar Labouta’s lab (College of Pharmacy, University of Manitoba). EV samples were diluted 1:75 with 0.22 µm filtered PBS. Five measurements were recorded for each sample with a dust cut-off set to 40. EV size was measured as an intensity averaged multimodal distribution using a scattering angle of 90°, and size bins were used to represent total size intensity within a given size range. Zeta potential analysis was performed using a Solvent-Resistant Electrode (NanoBrook) and BI-SCGO 4.5 ml cuvettes (NanoBrook). Smoluchowski formula was used to calculate zeta potential and final results were averaged irrespective of the charge (negative/positive) [13]. All measurements were performed in PBS (pH 7.4) at 25 °C.

### Statistical analyses

All data were analyzed using an unpaired Student’s t-test, except for size-distribution analysis for which we used a two-way repeated measures ANOVA. Multiple comparisions in the two-way ANOVA were corrected using Sidak’s post hoc test. Individual data points are plotted, with mean ± standard error of mean (SEM) shown as applicable. All graphs were created using GraphPad-Prism software (version 10.1.2, GraphPad, San Diego, CA, USA). Significance was set at p ≤ 0.05. A sample size of n = 8 biological replicates were used for each experiment.

## Results

### Chronic contractile activity alters EVs biophysical characteristics

To evaluate the effects of CCA on Skm-EVs, fully differentiated myotubes were electrically paced at 14 V (1 Hz) for 3 h/day for 4 consecutive days. After the last bout of contractile activity, myotubes were allowed to recover for 21h in sEV-depleted differentiation media. Conditioned media from both CON and CCA myotubes was then collected for EV isolation via dUC as illustrated in **Fig. 1A**. DLS analysis revealed that the mean size of CCA-EVs (132 nm) was ∼26% smaller than CON-EVs (178 nm) (p=0.013, n=8, **Fig. 1B**). A two-way repeated measures ANOVA on size distribution showed a significant main interaction effect between CON-EVs and CCA-EVs. Multiple comparision analysis further demonstrated that CCA-EVs had a 3.1-fold higher yield of 100-150 nm sized sEVs compared to CON-EVs (p=0.004, n=8, **Fig. 1C**). Conversely, CON-EVs exhibited a 100% increase in 200-250 nm EVs relative to CCA-EVs (p=0.003, n=8, **Fig. 1C**). These findings indicate that chronic contraction of myotubes induces a shift from lEVs to primarily sEVs. Lastly, zeta potential amalysis showed that CCA-EVs (−14.8 mV) had a 27% lower surface charge compared to CON-EVs (−11.6 mV) (p=0.024, n=8, **Fig. 1D**), suggesting potential differences in membrane composition and enhanced particle stability.

**Figure 1.**
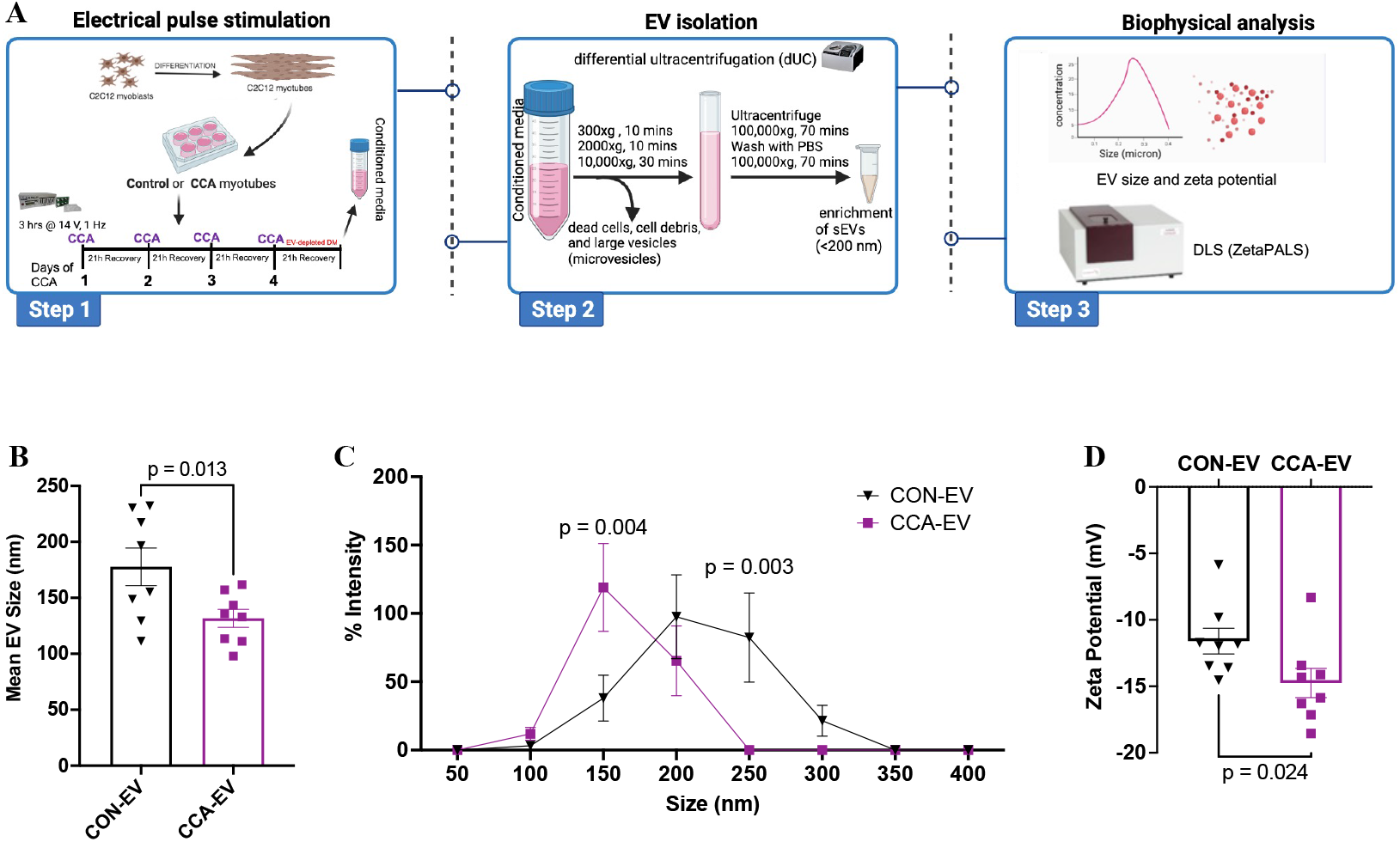
Effect of CCA on the biophysical properties of EVs. (**A)** C2C12 myoblasts were fully differentiated into myotubes (MTs) and divided into control (CON) and chronic contractile activity (CCA) groups. CCA-MTs were electrically paced (3h/day x 4 days at 14 V) using the IonOptix ECM to mimic chronic endurance exercise in vitro. Media was changed daily after each contractile activity bout, and cells were allowed to recover for 21 h. After the final bout of contractile activity, media was switched to differentiation media containing sEV-depleted horse serum in both CON and CCA myotubes, with an additional 21 h recovery period. Conditioned media from CON and CCA myotubes was then collected for EVs isoaltion via dUC. Isolated EVs were characterized by size and zeta potential using DLS (ZetaPALS). (**B)** Mean size, **(C)** Size distribution, and **(D)** Zeta potential in CCA-EVs vs CON EVs. Data were analyzed using an unpaired Student’s t-test in panels B and D, and by a two-way repeated measures ANOVA in panel C, with multiple comparisions corrected using Sidak’s post hoc test (n=8). Exact p values for significant results (p<0.05) are shown. Figure 1A created with BioRender.com.

## Discussion

In this study, we aimed to elucidate the effect of CCA on the biophysical properties of EVs released from C2C12 myotubes using DLS. Our results demonstrate that chronically stimulated C2C12 myotubes release smaller, more stable EVs than non-stimulated controls. To model chronic exercise in vitro, we used chronic electrical pulse stimulation of differentiated myotubes, a method previously shown by our group to effectively mimic chronic exercise-induced mitochondrial biogenesis [6]. In the present study, we observed a significant decrease in the mean size of CCA-EVs compared to CON-EVs. However, our previous findings using TRPS reported no difference in mean EV size but a higher concentration of CCA-EVs vs CON-EVs [6]. Additionally, size distribution analysis in the current study revealed that CCA-EVs were enriched with sEVs ranging from 100–150 nm, whereas CON-EVs were predominantly composed of 200–250 nm sized vesicles. Interestingly and in line with these findings, our previous TRPS analysis indicated that CCA-EVs were enriched with sEVs as well, with both 50–100 nm and 100–150 nm being higher compared to CON-EVs [6]. TRPS method uses single-particle analysis whereas DLS relies on estimating size from the average intensity of light scatter by EVs in solution. As such the former is more sensitive and can measure size accurately, whereas DLS can often overestimate size. It is encouraging that both methodologies indicate that CCA-EVs are enriched with particles ranging in size from 50 to 150 nm and underscores the robustness and reproducibility of results. The difference in size between CON-EVs and CCA-EVs found only with DLS in the current study is puzzling. It may be an artifact or a result of differential dispersant properties of the CON-EV samples.

Zeta potential is an important indicator of colloidal stability of particles in dispersed systems, such as suspensions, emulsions, and colloidal dispersions [14]. Zeta potential values further away from 0 indicate a greater degree of electrostatic repulsion and less tendency of aggregation or flocculation between particles [15]. Our result showed that EVs exhibited negative zeta potential values ranging from -5.9 to - 18.6 mV, consistent with previous reports [8], [16]. Notably, CCA-EVs had a significantly lower (more stable) zeta potential than CON-EVs. However, we did not observe a difference in zeta potential between groups in our previous findings using TRPS [6]. The biophysical properties of EVs may vary slightly depending on the characterization technique used, as reported in previous studies [8], [9]. Given that TRPS is a more sensitive, single-particle characterization method compared to DLS, it is better suited for analyzing polydisperse EV populations [8], [15]. DLS results, based on light scattering, inherently carry some degree of uncertainty, as larger particles scatter more light and can mask the presence of smaller vesicles. This methodological difference likely explains the discrepancies observed between DLS and TRPS measurements. Nonetheless, both techniques consistently showed that CCA-EVs were enriched in sEVs compared to CON-EVs. These findings align with previous studies reporting that endurance exercise training promotes the release of sEVs into circulation [17], [18], [19].

In summary, we demonstrated that CCA of murine myotubes induces the release of more smaller, and more stable Skm-EVs. These findings are consistent with previous reports indicating that chronic exercise alters EV biophysical characteristics. While our results align with prior TRPS-based analyses in some aspects, we also observed differences in size distribution and zeta potential, highlighting the influence of the measurement technique on EV characterization. Given the accessibility and ease of DLS, this method may serve as a useful alternative for assessing CCA-EVs in settings where more robust and sensitive techniques like TRPS are unavailable.

## Author Contributions

P.O.O. performed experiments in the current study, analyzed data, and created figures. P.O.O. and A.S. helped write and revise the manuscript. A.R.W., H.I.L. and J.W.G. provided technical and theoretical expertise to complete the work. All authors were involved in manuscript revisions. A.S. designed the project, and helped synthesize data, create figures, write, and edit the manuscript. A.S. is the corresponding author and directly supervised the project. All authors have read, edited and agreed to the published version of the manuscript.

## Conflicts of Interest

All other authors declare no conflict of interest. The funders had no role in the design of the study; in the collection, analyses, or interpretation of data; or in the writing of the manuscript.

Word count (not including title page, abstract, references, figure legends): 1890

